# Rapid, accurate and memory-frugal sub-gene-level quantification of probe-based, multiplexed single-cell data

**DOI:** 10.64898/2026.11.05.764594

**Authors:** Dongze He, Yuan Gao, Rob Patro

## Abstract

Probe-based, multiplexed single-cell assays profile multiple samples in parallel by sequencing known DNA probes tagged with both sample and cell barcodes. Most existing single-cell quantification tools were built for poly(A)-based assays and are not tailored to compact probe references or nested sample–cell barcodes.

We extend `simpleaf` to demultiplex and quantify probe-based single-cell libraries rapidly with a low memory footprint. In piscem, we index small probe panels in a cache-friendly hash dictionary for direct *k*-mer lookup, while a thread broker divides a shared thread budget between parallel FASTQ decompression and mapping. Downstream, we extend the RAD mapping-record format and `alevin-fry` with reusable, sample-scoped barcode correction tables (plans) and sort-free external-memory collation for *hierarchically* barcoded reads, and we preserve probe identities through UMI resolution.

Across five 10x Genomics Flex datasets spanning 44 samples, `simpleaf` had the shortest runtime and lowest peak memory use on every library, and produced gene-expression estimates, cell clusters, and marker genes concordant with cyto and `Cell Ranger`. Compared with cyto, `simpleaf` ran on average 2 times as fast and used roughly one-thirteenth as much memory. `simpleaf` also exports probe-level counts for sub-gene analysis. Using these counts, we uncover differential expression masked by gene-level aggregation, pinpoint the transcript groups behind gene-level changes, and resolve splicing differences across cell types.

## 1 Introduction

Single-cell RNA sequencing (scRNA-seq) profiles the transcriptomes of individual cells at scale [1]. Much of the progress in short-read, 3′-tagged scRNA-seq has come from changes in how RNA targets are captured and primed. Tang et al. [2] used anchored oligo(dT) primers to reverse-transcribe polyadenylated mRNA from isolated cells, which made poly(A) priming the standard approach for single-cell transcriptomics. 10x Genomics later combined oligo(dT)-functionalized gel beads, cell barcodes, and unique molecular identifiers (UMIs) with droplet microfluidics to profile thousands of cells in a single run [3]. Because oligo(dT) primers also anneal to internal adeninerich stretches, poly(A)-based libraries include reads from intronic, antisense, intergenic, and unannotated regions alongside 3′ transcript ends [4], [5]. More recently, probe-hybridization assays such as 10x Genomics Chromium Flex have seen rapid uptake [6]. In these assays, paired DNA probes hybridize to short target regions on transcripts and are ligated, barcoded, and amplified. Because hybridization requires only an intact 50-nucleotide target sequence instead of an intact poly(A) tail, these assays can profile fragmented RNA in fixed and archived tissues. Sequencing ligated probes also avoids the internal RNA priming and first-strand cDNA re-priming artifacts of oligo(dT) assays [7], and tagging probes with sample barcodes before pooling lets researchers process multiple samples in one library.

Widely used preprocessing tools such as `alevin-fry` [8] were built around poly(A)-captured cDNA from a single sample and do not take full advantage of how probe-based, multiplexed libraries are structured. Preprocessing these libraries differs in three ways: (i) biological reads come from a small, predefined panel of probe sequences instead of arbitrary 3′ transcript fragments; (ii) each probe has a known sequence, target gene, and splice-junction placement that determine which transcript isoforms it matches; and (iii) pooled samples share the same set of droplet cell barcodes, so a cell barcode identifies a cell only within its assigned sample.

Here we extend `simpleaf` [9], our unified preprocessing framework coordinating piscem [10] and `alevin-fry` [8], to process probe-based, multiplexed single-cell libraries end to end (Figure 1). To match the structure of these assays, we redesign each stage where poly(A)-oriented pipelines use unnecessary time or memory (Section 2). During mapping, piscem queries a compact hash dictionary tailored to small probe panels while dynamically splitting threads between parallel FASTQ decompression and read mapping, and writes probe assignments, sample and cell barcodes, and UMIs to Reduced Alignment Data (RAD) v2 records, a compact binary format for passing mapped reads to `alevin-fry`. Downstream, `alevin-fry` precomputes sample-scoped barcode corrections once per distinct barcode, groups (collates) records by sample–cell identity on disk with-out a comparison sort or per-bucket temporary files, and retains probe identities through UMI resolution (collapsing duplicate reads per molecule) to output both gene and probe count matrices. Across five 10x Genomics Flex benchmarks, `simpleaf` cuts both runtime and peak memory relative to existing tools while matching their gene-expression estimates and down-stream clustering results. Retaining individual probe counts also supports sub-gene analyses of differential expression, isoform usage, and splicing that gene-level totals obscure.

**Figure 1.**
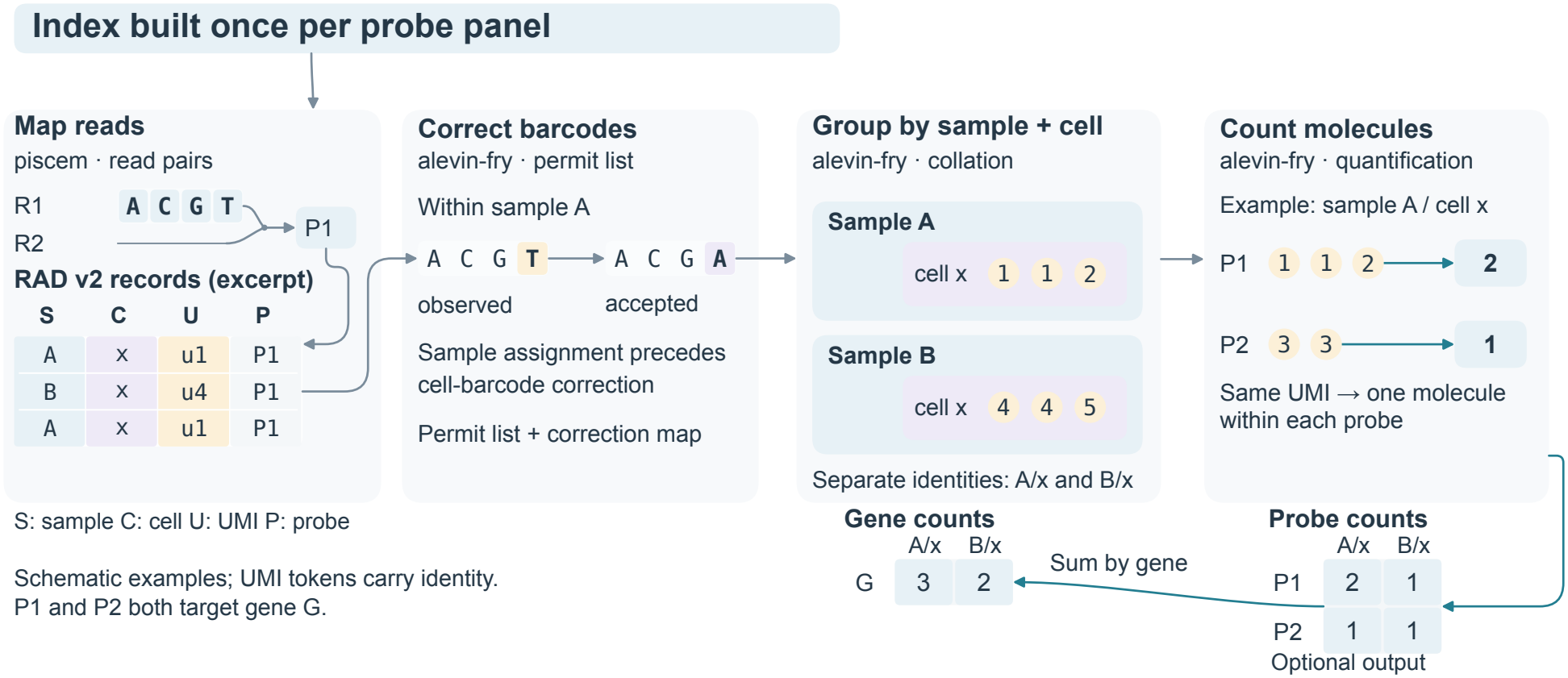
Processing probe-based, multiplexed single-cell data with `simpleaf`. A probe index is built once per probe panel and reused across experiments. piscem maps read pairs and writes probe assignments, sample and cell barcodes, and UMIs to RAD v2. `alevin-fry` generates a permit list and correction maps, corrects cell barcodes within assigned samples, and groups records by sample–cell identity. Cell barcode x in sample A remains distinct from x in sample B. UMI resolution produces molecule counts within each probe; repeated tokens denote identical UMIs. Counts for probes P1 and P2 sum to gene G, and probe counts can also be output directly. Sequences and counts are schematic, and only excerpts of records and groups are shown. Read-field positions and chemistry-specific structure are omitted.

## 1 Methods

Quantifying a probe-based, multiplexed library requires mapping short reads against a probe panel, correcting nested sample and cell barcodes, and grouping the mapped records by sample and cell for UMI resolution (Figure 1). A typical Flex experiment contains billions of reads, yet only tens of thousands of probes and a few hundred thousand distinct barcodes. To exploit this asymmetry, we shift work off the per-read path onto the probe panel and the set of distinct barcodes, pairing direct *k*-mer hashing (Section 2.1) and dynamically balanced decompression (Section 2.2) with precomputed barcode correction (Section 2.3), sort-free collation (Section 2.4), and probe-level UMI resolution (Section 2.5).

### 2.1 A dictionary specialized for probe panels

When mapping to a full transcriptome, piscem uses sshash [11], [12], which saves memory across tens of millions of *k*-mers by indexing minimizers with a minimal perfect hash and scanning candidate strings in a compact *k*-mer representation (a spectrum-preserving string set). A Flex probe panel, however, contains fewer than two million canonical *k*-mers (the lexicographically smaller of each *k*-mer and its reverse complement), which is small enough to trade extra bytes per key for direct lookup. Our `tiny-dict` back-end therefore stores every canonical *k*-mer explicitly in a hashbrown open-addressing hash table (based on SwissTable [13]), so lookups need neither minimizer extraction nor string scanning (Figure 2)

Because a *k*-mer can appear in more than one probe, `tiny-dict` retains `piscem’s` two-step indirection from *k*-mers to non-branching segments (unitigs) of the compacted probe de Bruijn graph and from segments to probes, keeping one hash entry per *k*-mer while reusing `piscem’s` downstream mapping engine. Each table entry contains the 64-bit canonical *k*-mer key alongside a packed segment location (segment identifier, offset, and orientation), so the mapper verifies exact membership and reads the match coordinates in a single table lookup without a separate value-array access. We then resolve segment coordinates to probe coordinates through a segment-occurrence table, where each entry gives the probe occurrences for one segment. Because probes are designed to be distinct in sequence, nearly every segment occurs in only one probe; we pack those single-probe coordinates inline in the segment-occurrence entry and place only multiprobe segments in an overflow list. On almost all positive queries, piscem therefore avoids chasing and decoding an external occurrence list, and lookups stay cache-local.

**Figure 2.**
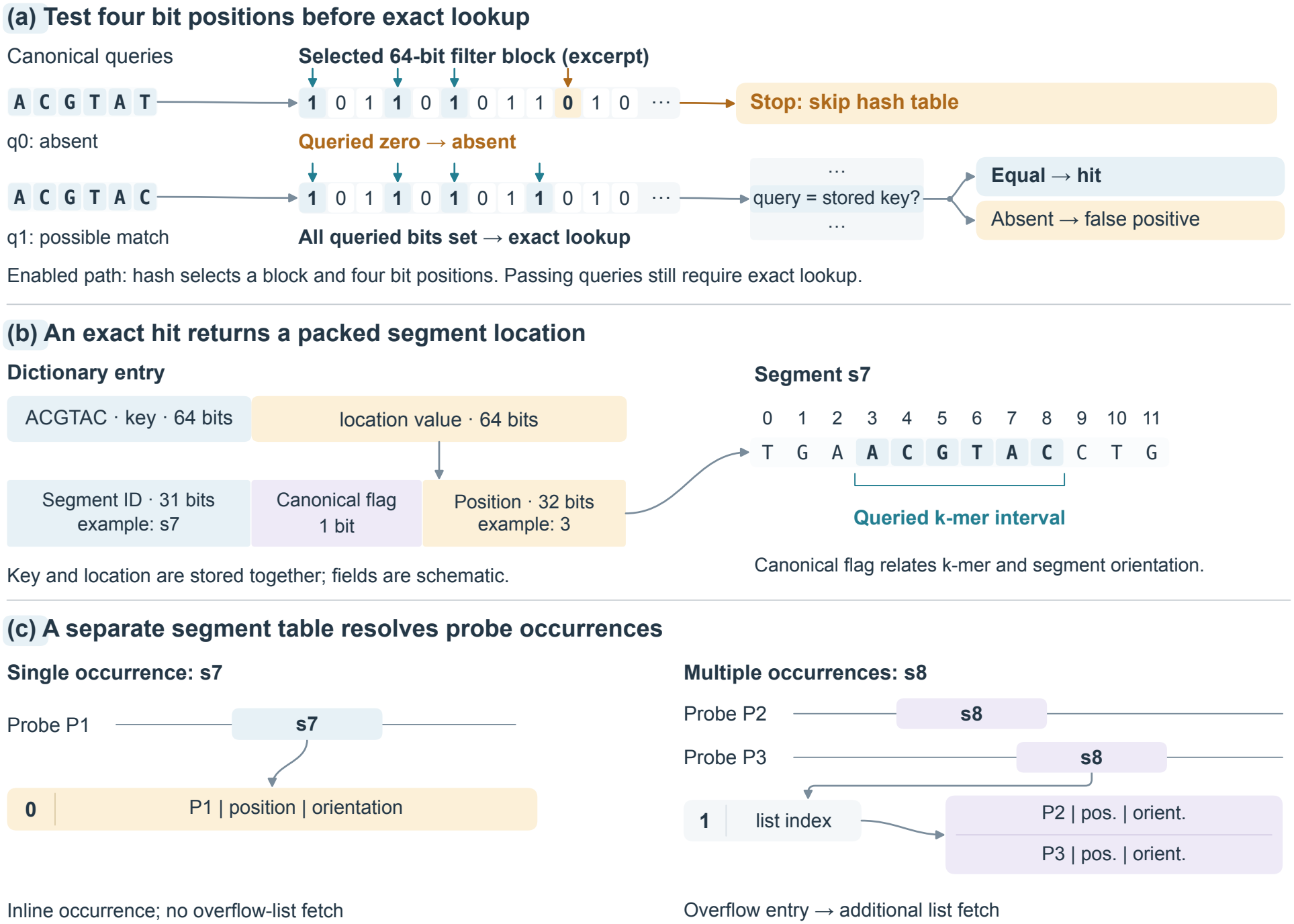
Lookup paths in the small-reference index. (a) When enabled after size and sampled-miss-rate checks, the mapper tests four hash-derived positions in one 64-bit block of the register-blocked Bloom filter. A queried zero rules out membership; if all four bits are set, the mapper performs an exact dictionary lookup to distinguish a true hit from a false positive. Arrows mark tested positions in two illustrative queries; positions may coincide in practice. (b) On an exact hit, the lookup returns a packed segment location stored beside the canonical *k*-mer key. The value contains a 31-bit segment identifier, a 1-bit canonical-orientation flag, and a 32-bit within-segment position. In the shortened sequence, the queried interval begins at zero-based position 3 of segment s7. (c) Each entry in the segment-occurrence table contains either a single inline probe occurrence (high bit 0) or an overflow-list index (high bit 1), illustrated by s8 shared between two probes. Fields and sequences are schematic; field widths are not drawn to bit scale, and occurrence-field widths depend on the reference.

When sequencing reads extend past the 50-bp probe into adapter sequence, their trailing *k*-mers are absent from the panel and trigger repeated dictionary misses. To reject absent *k*-mers cheaply, tiny-dict places a register-blocked Bloom filter [14] in front of the hash table and tests four hash-derived bit positions within a single 64-bit word. Unlike a standard Bloom filter that scatters bit tests across multiple cache lines, this layout lets the mapper inspect at most one cache line per filter check. Keys with all four bits set are still verified in the hash table, so filter false positives add only an occasional lookup and never alter mapping assignments. Because the filter adds a bit check to true hits, we cap its size at 2 MiB so it stays cache-resident and enable it only when sampled queries show a sufficiently high miss rate.

Across the three probe-panel indexes used in this study (two human panel versions and one mouse panel), each index contained 1.71–1.77 million distinct canonical *k*-mers, lookup structures accounted for 37.46–37.52 MiB of requested heap allocations (including the 2 MiB prefilter), and 99.63–99.78% of segments had inline occurrence entries (Supp. Table 4).

### 2.2 Balancing decompression and mapping

Fast probe lookups let piscem consume reads faster than a single gzip stream can decompress them. Parallel decompression removes that input bottleneck, but a fixed thread split between decoding and mapping is fragile because compression ratios, read lengths, hit rates, and storage speeds vary across runs. Too few decoders leave mappers waiting on input, whereas too many take cores away from mapping and inflate inmemory read buffers. Because standard gzip blocks depend sequentially on earlier deflate dictionaries, we port the speculative decoding and dictionary-back-filling algorithm of `rapidgzip` [15] (which decodes file chunks in parallel and resolves backward references once preceding chunks complete), pair it with paraseq FASTQ parsing [16], and add a thread broker that dynamically divides a shared budget of *N* threads between decoding and mapping.

Following rate-based dataflow control [17], the broker separates active CPU computation from time spent waiting on queues. Over steady-state windows where the amount of buffered work barely changes, the decoder and mapper process roughly the same number of records. The broker measures their aggregate active thread times *B*_*d*_ for decoding and *B*_*M*_ for mapping (excluding time blocked on the input source or output queue), so the ratio *B*_*d*_/*B*_*M*_ reflects the relative perrecord computational cost of the two stages. To match their service rates within *N* total threads, it sets the idealized target allocation as

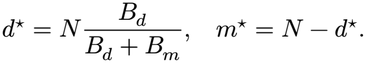

In practice, the broker smooths these measurements, ignores startup and resize transients, rounds and bounds *d*^⋆^, and stops adding decoders when input cannot be read faster. When shifting threads, it shrinks one pool before expanding the other so total concurrency stays within *N*, and it rolls back any adjustment followed by a drop in throughput. Beyond Flex preprocessing, this approach is useful in other high-throughput genomics pipelines where decompression and parsing would otherwise leave worker threads idle.

### 2.3 Correcting identities before streaming records

Once piscem writes mapped reads to a RAD file, `alevin-fry` detects and corrects barcodes, collates records, and resolves UMIs. In a multiplexed library, all pooled samples share the same set of droplet cell barcodes, so a cell barcode identifies a cell only within its sample of origin. Moreover, the set of real cells and their exact-match barcode counts differ from sample to sample, so sample assignment must precede cell-barcode correction. For an observed sample barcode *s* and cell barcode b, let S(*s*) denote the accepted sample correction and C_S(*s*)_(b) the cell correction within sample S(*s*). Each cell is identified by the ordered pair (S(*s*), C_S(*s*)_(b)), so identical cell-barcode sequences in different samples remain separate and UMIs are resolved independently within each pair. While Flex uses two barcode levels, we designed the RAD v2 schema and correction pipeline to generalize to deeper barcode hierarchies.

In RAD v2, barcode roles and hierarchy levels are recorded once in the file prelude (the header meta-data at the start of the file), so downstream tools can reconstruct composite identities directly from the schema (Supp. Figure 1). Because billions of reads repeat a much smaller set of observed barcodes, `alevin-fry` separates barcode resolution from record streaming. During its first pass (permit-list generation), `alevin-fry` determines the set of accepted cell barcodes (the permit list) for each sample and precomputes a sample-scoped *correction plan*, a compact serialized table containing each distinct observed barcode’s final decision within its sample (its accepted target b^′^ or an uncorrectable flag), which collation reuses via direct lookups instead of repeating neighborhood searches per read.

To build the permit list and correction plan, `alevin-fry` first routes reads with unambiguous sample barcodes immediately through a precomputed lookup table. When an observed sample barcode lies one mismatch from multiple valid sample barcodes so tie-breaking requires exact-match sample frequencies, `alevin-fry` spools compact sample–cell barcode pairs and replays them once those counts are tallied. To avoid per-read synchronization while counting cell barcodes, workers tally counts in bounded local maps and periodically merge them into per-sample histograms.

Within each sample, permitted cell barcodes are stored as sorted, two-bit-encoded integers indexed by a prefix-offset directory containing the start offset of each short barcode prefix in the sorted array. To test an observed barcode, `alevin-fry` uses its prefix to select a narrow contiguous slice of the array for binary search and generates one-substitution neighbors directly via bit operations on the two-bit integer. Each neighbor is looked up under its own prefix, including substitutions that alter the prefix bases, so every check stays confined to a narrow slice instead of scanning the full permit list. Exact matches take precedence; for one-mismatch neighbors, `alevin-fry` either requires a single matching target (the unique-target policy, discarding ties) or breaks ties in favor of the neighbor with higher exact-match support (the frequency-based policy, using priors fixed from exact-match counts). Because `alevin-fry` writes one resolved decision per distinct observed cell barcode into the sample-scoped correction plan, neighborhood search scales with distinct barcodes instead of read count.

To prepare for external-memory collation (Section 2.4), `alevin-fry` uses the per-cell read counts to pack corrected sample–cell groups into memorysized *logical buckets* (batches of cells that can be collated independently within RAM without splitting any cell’s records) and compiles the saved correction plan into a sample-routing table plus per-sample cell tables. During collation, `alevin-fry` maps each observed sample–cell pair through sample routing and a single cell-barcode lookup to obtain both the corrected cell barcode and its target bucket (packed into one integer when narrow enough, or stored in a wider entry otherwise). Every retained record with the same corrected (S(*s*), C_S(*s*)_(b)) pair is routed to the same logical bucket so all records for a cell can be gathered together for independent UMI resolution.

### 2.4 Collation without a comparison sort

UMI resolution requires all records for a corrected sample–cell pair to be stored contiguously, yet the full RAD file can be far larger than available RAM. We collate records in external memory with a two-phase scatter–gather pass guided by the bucket assignments from permit-list generation, where scatter workers partition incoming records into logical buckets on disk and gather workers later read each bucket back into memory to group records by cell. Opening a separate temporary file per bucket scales poorly on large libraries because hundreds of buckets can exhaust file descriptors and fragment disk writes. Instead, each scatter worker appends to a single private temporary (spool) file and logs the byte ranges (extents) written for each logical bucket, so the number of temporary files equals the worker count.

Because UMI resolution requires only that records sharing a sample–cell identity be contiguous (not lexicographically ordered), gather workers group each logical bucket in two linear passes over its spooled extents instead of running a comparison sort. In the first pass, a gather worker reads only record headers to extract each record’s corrected sample–cell key and byte length, sums the bytes needed per group, and tags each record in memory with a 32-bit group index. From those totals, the worker computes cumulative byte offsets (prefix sums) to lay out disjoint, contiguous output slices for every group. In the second pass, the worker uses the saved group indices to copy raw record bytes directly into each group’s next free offset without re-hashing multi-barcode keys or decoding alignment payloads. Once the pass completes, all records for each corrected sample–cell identity occupy a single contiguous block.

To keep memory bounded, `alevin-fry` sizes spool buffers and gather concurrency from the bucket-size estimates and a configurable working-memory target, and frees the scatter lookup tables before allocating gather output buffers so the two working sets never overlap. This target applies to collation buffers and does not hard-cap total process resident set size (RSS), which also includes correction metadata, allocator overhead, and auxiliary buffers. Storage layouts and lookup details are given in Section S2.

### 2.5 Retaining probe identity through UMI resolution

Because piscem writes matched probe identifiers instead of gene identifiers to the RAD file, `alevin-fry` can deduplicate UMIs at either the gene or probe level from the same barcode-corrected, collated records without re-running mapping or collation. The user selects the resolution through a reference-to-feature map by either grouping all probes of a gene into one feature for gene-level counts or leaving each probe as its own feature for probe-level counts.

Within each corrected sample–cell group, `alevin-fry` resolves UMIs at the chosen feature level. Under the cr-like strategy (matching `Cell Ranger’s` rule), each UMI is assigned to the feature with the strongest read support and discarded on an unresolved tie, and one-mismatch UMI error correction operates within that same feature scope. Consequently, reads from different probes of the same gene can support a gene-level UMI assignment even when probe-level support is tied, and one-mismatch UMIs on different probes of a gene are collapsed together only during gene-level resolution. Because each probe hybridizes independently to a distinct locus on a transcript, `simpleaf multiplex-quant` resolves UMIs per probe by default and sums each gene’s probe counts to form the gene matrix (in practice, summed probe counts and direct gene-level UMI counts agree to within 0.23%; Section S3). We use this default throughout our benchmarks and analyses.

## 3 Results

We evaluated `simpleaf` across Flex v1 and v2 libraries with different multiplexing designs, comparing its runtime, memory footprint, and gene-level concordance against cyto [18] and Cell Ranger [3] before examining what probe-level counts add beyond gene totals.

### 3.1 simpleaf is fastest and most memory-frugal across the benchmarked libraries

We benchmarked `simpleaf, cyto`, and Cell Ranger on five Flex datasets spanning human and mouse tissues, Flex v1 and v2 chemistries, 4- and 16-plex designs, and sequencing depths from hundreds of millions to 11 billion reads (Section 5.3). All runs executed in Docker containers under matched CPU and memory limits (Section 5.1; Figure 3).

**Figure 3.**
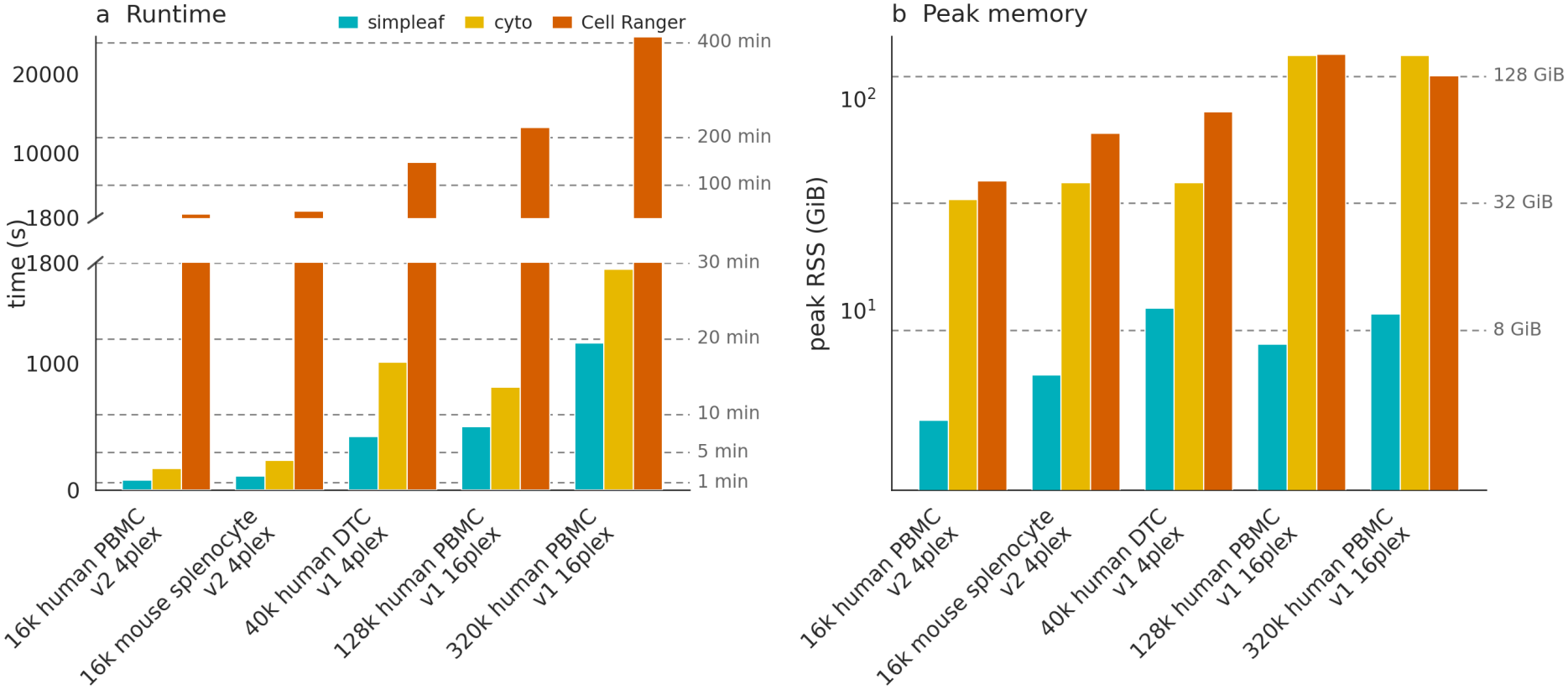
Runtime and peak resident set size for `simpleaf` under one-mismatch UMI collapse, `cyto` and `Cell Ranger`, ordered by read count. **a**, runtime. **b**, peak resident set size of the whole process tree. The runtime axis is broken at 1,800 s, with the two segments on different linear scales; the memory axis is logarithmic.

`simpleaf` had the shortest runtime and lowest peak memory on every library. cyto took an average of 2.21 times as long on the 4-plex libraries and 1.56 times as long on the 16-plex libraries, while using 3.9 to 23.3 times as much memory (12.6 times on average); Cell Ranger took 20.9 to 28.8 times as long (24.1 times on average) and used 8.5 to 23.5 times as much memory (14.6 times on average).

Runtime in `simpleaf` is dominated by read mapping and scales linearly with read count. Across 783 million to 11.0 billion mapped reads, throughput remained steady at 8.6–9.7 million reads per second for `simpleaf`, compared with 3.6–6.3 million for cyto. Sustaining that mapping rate across compressed FASTQ files depends on parallel gzip decoding (Section 2.2), which `simpleaf` enables by default on Flex data whenever at least four threads are available per input gzip file (when fewer threads per file are available, reading multiple files concurrently already parallelizes decompression; configurable via --thread-policy).

Peak memory in `simpleaf` stayed between 3.0 and 10.2 GiB across all five libraries, determined by quantification on the smallest datasets and by mapping on the larger ones. Because tiny-dict requires about 37.5 MiB for the probe index, peak memory during mapping comes mainly from in-flight reads buffered between FASTQ decoders and mapping threads. The thread broker sizes decoding throughput to keep the fixed pool of mapping threads fed, so buffer memory scales with thread count rather than library depth. Processing 11 billion mapped reads (709 GiB of compressed FASTQ input) required 9.6 GiB of RAM, and a 14-fold increase in read count raised peak memory only 3.4-fold. Across the five libraries, `simpleaf`’s average peak memory (6.9 GiB) was 8.0% of cyto’s (86.7 GiB) and 7.1% of Cell Ranger’s (97.5 GiB).

### 3.2 Gene counts are concordant between tools

Across all five libraries (44 samples), we compared gene counts pairwise among `simpleaf`, cyto, and Cell Ranger on the high-confidence cells called by Cell Ranger (Section 5.4.4). Predefined probe sequences make counts directly comparable across tools, and multimapping is rare because probes are chosen to be distant in sequence space. All three tools agreed closely across the 44 samples, with mean percell Spearman correlations of 0.966 to 0.997, count concordance of 97.8% to 99.6%, and mean absolute relative differences of 0.002 to 0.024. In the 16k mouse splenocyte Flex v2 library, for instance, `simpleaf` and cyto reported 1.02% and 1.20% fewer total UMIs than Cell Ranger on shared cells and genes. In a read-level audit of `simpleaf`, we traced most of this difference to Read 1 (R1) barcode-geometry parsing and a smaller fraction to Read 2 (R2) probe matching (Section S1).

On R1 in Flex v2, a variable-length spacer sits between the cell-barcode/UMI prefix and a fixed 12-nucleotide anchor sequence preceding the sample barcode, so the sample barcode’s offset shifts from read to read. cyto searches directly for a sample barcode at its expected offset (allowing one unambiguous substitution) and, on a miss, steps outward up to five bases in either direction without checking the upstream anchor [19]. `simpleaf` and `Cell Ranger` instead align the 12-nucleotide upstream anchor allowing up to one mismatch. Under its default Flex v2 geometry, `simpleaf` tests five candidate anchor offsets (spacers of 0–4 nucleotides) and extracts the sample barcode immediately downstream of the best-matching anchor. `Cell Ranger` can additionally recover sample barcodes at five predefined offsets when no anchor matches, and checks anchor offsets outside `simpleaf`’s window if the resulting barcode matches the known sample-barcode whitelist exactly [20]. These edge cases affect few molecules. In the mouse splenocyte library, for example, reads with an anchor one nucleotide upstream of `simpleaf`’s window accounted for 1.64% of read pairs and were retained by Cell Ranger but excluded by `simpleaf` (Section S1).

On R2 (the 50-nucleotide probe read), minor mapping differences stem from how each tool tolerates sequencing errors. By default, `simpleaf` queries exact 19-mers, so sensitivity depends on error placement as well as error count. Because a 50-base probe spans 32 overlapping 19-mers, two well-spaced substitutions can disrupt every 19-mer and leave a read unmapped, whereas two substitutions near the ends leave intact 19-mers in between. `Cell Ranger` and cyto instead split each probe into two 25-base halves and initially allow up to one substitution per half. When only one half matches, both tools use that half to select candidate probes and check the second half under different rescue rules. `Cell Ranger` computes a bounded edit distance permitting substitutions, insertions, and deletions subject to alignment-score thresholds [21], whereas cyto permits up to three substitutions across the full probe provided the assignment is unambiguous [19].

### 3.3 Biological interpretations at the gene level are concordant between tools

To test whether these small count differences affect downstream interpretation, we ran a standard single-cell clustering and marker-discovery workflow [22] independently on each tool’s count matrix. Here we highlight the 128k 16-plex human peripheral blood mononuclear cell (PBMC) library (7,190 to 8,490 cells per sample), with results for the remaining four datasets summarized in Supp. Table 5 (pooled UMAP embeddings in Supp. Figure 2). After tuning the Leiden resolution so all three tools produced the same number of clusters per sample (22 on average), pair-wise cluster homogeneity and completeness across the 48 comparisons (3 tool pairs across 16 samples) averaged 0.936 and 0.938, respectively.

Cluster marker genes agreed closely as well. Among the top 50 markers per cluster, an average of 45 genes matched between Cell Ranger and cyto (mean Jaccard index 0.851) and between Cell Ranger and `simpleaf` (mean Jaccard index 0.868).

### 3.4 simpleaf reveals biology buried in the gene counts

Unlike poly(A)-primed reads, whose splicing status and isoform compatibility must be inferred computationally [8], [23], each Flex probe has a fixed target sequence with known splice-junction placement and transcript compatibility. Exporting a probe-by-cell count matrix therefore makes sub-gene expression variation directly testable [24]. Using CellTypist cell-type annotations ([25]; Section 5.4.5), we compared gene- and probe-level expression across immune populations in the 320k human PBMC library. We assessed differential expression separately at the gene and probe levels by paired pseudobulk analysis with PyDESeq2 [26], using Wald tests with Benjamini– Hochberg correction (Section 5.4.6).

Summing probe counts into a single gene total can hide significant regulation of individual probes when an unchanging high-abundance probe dominates the sum or when probes shift in opposite directions (Figure 4, panel a). In naive B cells, *ZBTB16* showed no gene-level differential expression (fold change 1.02; adjusted p = 0.85) even though one of its three probes changed significantly, because the highest-count probe remained nearly constant and diluted the signal. In memory B cells, *FOSB* had opposing probe-level changes, where one probe decreased by 10% and another increased by 37%, with consistent directions across 10 and 11 donors, respectively. Both probe-level shifts were statistically significant, whereas their sum showed no significant gene-level change (fold change 1.06; adjusted p = 0.14).

**Figure 4.**
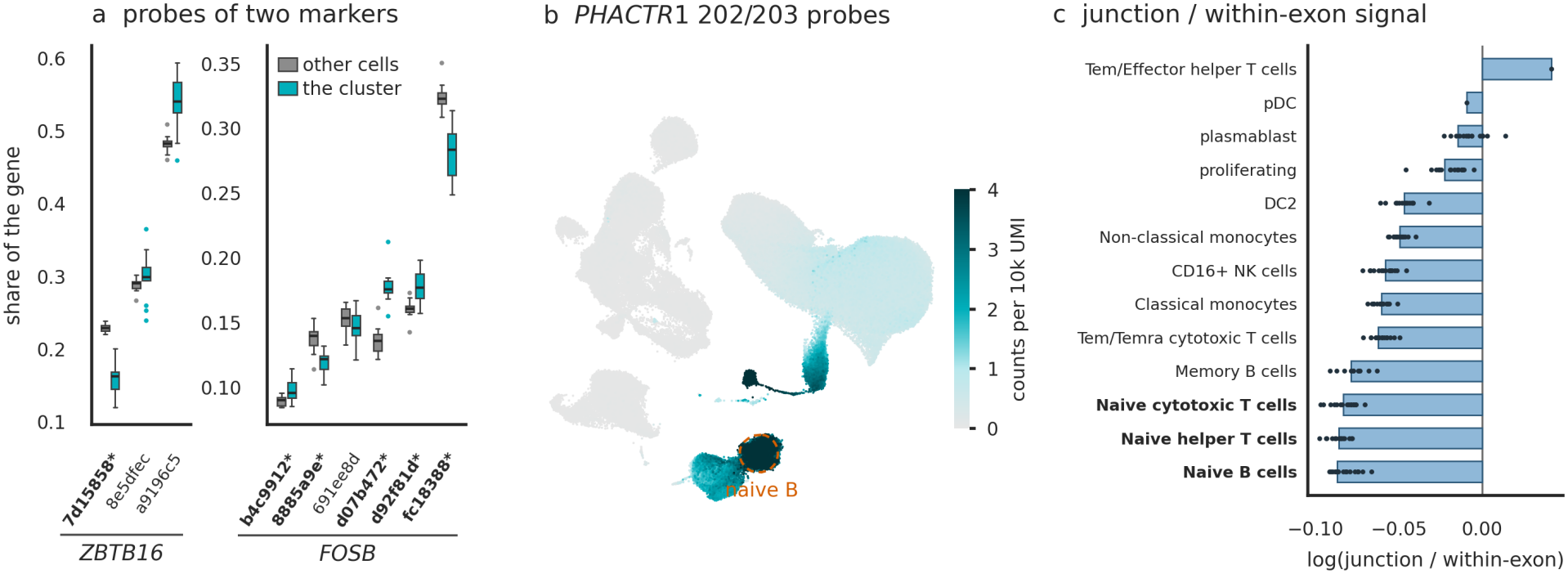
Sub-gene expression patterns in the 320k human PBMC library. **a**, Cell-type marker genes without significant gene-level differential expression (DE) because a DE probe is diluted by higher-count unchanged probes (*ZBTB16*) or because individual probes change in opposite directions and cancel out (*FOSB*). **b**, *PHACTR1* probes targeting isoforms *PHACTR1*−202 and −203 are enriched in naive B cells, whereas probes targeting other *PHACTR1* isoforms are rarely detected. **c**, Log ratio of junction-spanning to within-exon probe counts across cell types; naive immune populations have lower ratios than mature and proliferating populations.

Even when a gene changes strongly overall, we can use individual probe counts to narrow down which transcript isoforms account for that change because different probes for the same gene match distinct subsets of isoforms. In naive B cells, *PHACTR1* expression was 7.5-fold higher at the gene level than in other cells (Figure 4, panel b). That increase came entirely from the three probes matching isoforms *PHACTR1*−202 and *PHACTR1*−203, whose counts rose 6.1-, 6.2-, and 11.4-fold; the remaining three probes, which match other *PHACTR1* isoforms, accounted for just 52 UMIs across the entire library and zero in naive B cells.

We can also separate spliced mRNA from total (spliced plus unspliced) RNA by comparing junction-spanning probes against within-exon probes. A junction-spanning probe hybridizes across an exon–exon boundary and therefore captures only spliced tran-scripts, whereas a within-exon probe captures both spliced mRNA and unspliced nascent or intron-retaining RNA. Restricting each gene to junction-spanning and within-exon probes that match the same set of isoforms (Section 5.4.7), we observed in the 320k human PBMC library (Figure 4, panel c) that naive B and T cells had a consistently lower ratio of junction-spanning to within-exon probe counts than their mature counterparts and proliferating cells, consistent with splicing differences between naive and mature lymphocyte states.

## 4 Discussion

Although multiplexed, probe-based assays add sample demultiplexing and a probe-to-gene layer to single-cell preprocessing, their small reference and bounded barcode space let us simplify the most expensive per-read steps. Matching each stage to this scale asymmetry, `piscem` replaces compressed transcriptome indexing with direct hash lookups and balances parallel FASTQ decompression against mapping, while `alevin-fry` precomputes sample-scoped barcode corrections once per distinct barcode and collates records on disk without a comparison sort or per-bucket temporary files. Across all five Flex benchmarks, `simpleaf` outperformed cyto and Cell Ranger in runtime and memory while matching their gene counts, clusters, and markers.

Two design boundaries and evaluation caveats are worth noting. First, because we benchmarked the complete workflow end to end, separate ablation experiments would be needed to isolate the individual speedup of each algorithmic component. Second, tiny-dict’s direct hash table is specialized for small references of up to a few million *k*-mers, so piscem retains sshash for full transcriptomes and genomes. Conversely, our dynamic decompression-to-mapping thread broker and the hierarchical RAD v2 barcode schema are assay-agnostic and apply directly to deeper combinatorial barcoding protocols and other high-throughput sequencing pipelines bottlenecked by gzip input.

Finally, retaining probe identities through UMI resolution adds minimal quantification overhead while preserving sub-gene signal lost under gene-level aggregation. Using the probe count matrix from human PBMCs, we uncovered masked regulation, pinpointed the isoforms behind a gene-level shift, and resolved splicing differences between naive and mature lymphocytes from standard outputs. Because fixed probe panels hybridize only to predefined target sequences, they do not capture novel exons or unannotated splice junctions outside the designed set; for covered loci, however, exporting probe counts makes sub-gene quantification immediately accessible alongside routine gene-level analysis.

## 5 Experimental setup and analysis

### 5.1 Environment setup

All benchmarks were performed in Docker containers on an AWS i4i instance with an Intel Xeon Platinum 8375C processor, 64 vCPUs, and 495 GiB of memory, running Ubuntu 24.04. For the comparison across tools, each container was limited to 32 CPUs and 300 GiB of memory, with input files staged on local NVMe storage. We used `simpleaf` 0.31.0 [9], `alevin-fry` 0.19.0 [8], `piscem` 0.24.0 [10], cyto 0.4.6 [18], and `Cell Ranger` 10.1.0. The Rust-based tools were compiled from source in release mode with native CPU optimizations; `Cell Ranger` was used as distributed.

### 5.2 Datasets

We analyzed five Flex datasets: a 128k human PBMC TotalSeq-C 16-plex library, a 320k human PBMC sub-pool 16-plex GEM-X library, a 40k human dissociated tumor cell (DTC) library from three donors, a Flex v2 16k human PBMC TotalSeq-C 4-plex library, and a Flex v2 16k mouse splenocyte 4-plex library. Dataset download pages are provided in Supp. Table 1.

### 5.3 Benchmarking configuration

Gene-expression FASTQ files were processed in Docker containers by calling the `simpleaf multiplex-quant, cyto workflow gex,` and `cellranger multi` programs, with the official probe sets and barcode lists provided by 10x Genomics for each chemistry. Each program used its default Flex analysis settings, subject to the resource limits described in Section 5.1. We measured wall-clock run-time and peak RSS across the full process tree for each run.

For `simpleaf` and cyto, each dataset was processed four times, with the first run discarded as a warmup. Input files were evicted from the filesystem cache between runs. We reported the minimum runtime and maximum peak RSS across the three measured runs. `Cell Ranger` was run once per dataset without a warmup because repeated runs were prohibitively time-consuming.

### 5.4 Data analysis

We analyzed the count matrices generated by the three tools with Scanpy [27], following published recommendations for single-cell RNA-seq analysis [22].

#### 5.4.1 Cell selection and data loading

Cell calling is not evaluated in this work; instead, we used `Cell Ranger’s` filtered cell barcodes to define the initial cell set for all three tools. Gene-expression matrices from `Cell Ranger,` cyto, and `simpleaf` were loaded into Scanpy as AnnData objects, and cell and gene identifiers were aligned across tools so that all comparisons used identical cell and gene sets. Sub-sequent quality control (Section 5.4.2) and clustering (Section 5.4.3) were run independently on each tool’s matrix.

#### 5.4.2 Quality control and doublet removal

Low-quality cells were filtered by median absolute deviation (MAD) thresholds of five MADs for general quality-control metrics and three MADs for mitochondrial count fraction; cells exceeding 8% mitochondrial counts were also removed. We corrected ambient RNA contamination with SoupX [28] and excluded genes detected in fewer than 20 cells.

Doublets were called with scDblFinder [29], retained through clustering and cell-type annotation, and removed prior to probe-level differential expression testing.

#### 5.4.3 Normalization, feature selection, and clus tering

Counts were normalized by pooling-based size-factor estimation in scran [30]. We selected the 4,000 most informative genes by deviance in scry [31], ran principal-component analysis on those genes, built a nearest-neighbor graph on the first 30 principal components, and clustered cells with the Leiden algorithm [32].

To compare clusterings at matched granularity, we tuned the Leiden resolution on each count matrix to match the cluster count produced on the Cell Ranger matrix at resolution 1.0.

#### 5.4.4 Concordance between tools

We compared gene-expression estimates on the shared cell and gene sets using mean per-cell Spearman correlation, count concordance, and mean absolute relative difference as defined previously [8]. Cluster partitions were compared using adjusted Rand index, normalized mutual information (normalized by the arithmetic mean of the two partition entropies), adjusted mutual information, V-measure, Fowlkes– Mallows index, normalized variation of information, homogeneity, and completeness.

To assess marker-gene agreement, we ranked cluster markers by a Wilcoxon rank-sum test, paired clusters across tools by maximum-overlap (Hungarian) matching [33], and computed the number of shared genes and Jaccard index among the top 50 markers per cluster.

#### 5.4.5 Celltype annotation

We annotated cell types on the `simpleaf` gene-expression matrix with CellTypist 1.7.1 [25] (`Immune_All_Low` model) using majority voting over Leiden clusters.

#### 5.4.6 Probe annotation and differential expres sion

To determine each probe’s transcript compatibility and position relative to splice junctions, we matched its 50-nucleotide sequence exactly (in both orientations) against annotated transcripts, using the 10x Genomics GRCh38-2024-A reference for the human panels. Probes matching wholly within a single exon were classified as within-exon probes, and probes crossing an exon–exon boundary as junction-spanning probes; the transcripts containing a probe’s sequence formed its compatible transcript set.

For each cell type and donor, we summed gene and probe counts into paired pseudobulks for the target cell type versus all remaining cells and tested for differential expression in PyDESeq2 [26] (∼ donor + grp). Gene-level and probe-level models were fit separately, each with its own median-of-ratios size factors and dispersion estimates, and evaluated by Wald tests with Benjamini–Hochberg correction. Probe testing was restricted to probes with at least 10 UMIs in both groups. A cell type was tested when it appeared in at least three donors with at least 100 cells in total. A donor was included in a pseudobulk pair only when at least 25 cells of the target cell type were observed in that donor, and each test required at least three qualifying donors. At the probe level, Benjamini–Hochberg correction was applied across the tested probes. Fold changes were reported without shrinkage, and all tests used PyDESeq2 0.5.4.

We flagged a probe-level change as masked by gene-level aggregation when the probe had an adjusted p-value below 0.05 while its parent gene had an adjusted p-value of at least 0.05, and used probe–transcript compatibility sets to identify the isoform groups associated with differentially expressed probes.

#### 5.4.7 Analysis of splicing information in probe based assays

To compare spliced versus total RNA signal across cell types without confounding from alternative isoform usage, we restricted each gene to junction-spanning and within-exon probes that shared identical compatible transcript sets. For each gene, we divided junction-spanning and within-exon count sums by their respective probe counts so unequal probe numbers would not bias the comparison, and summarized the log ratio across genes and donors per cell type. We retained genes with defined ratios in every cell type represented by at least three donors so rare single-donor cell types would not overly restrict the shared gene set.

## 6 Code availability

`simpleaf` is implemented in Rust and distributed under the BSD 3-clause license at github.com/ COMBINE-lab/`simpleaf`. Benchmark and analysis scripts and input-reference requirements are documented at https://github.com/COMBINE-lab/fast-flex-processing-paper-scripts.

### Declarations

R.P. is a co-founder of Ocean Genomics Inc.

## Acknowledgements

This work was supported by the US National Institutes of Health (R01HG009937) and by grants 2022-311195 and 2024-342821 from the Chan Zuckerberg Initiative DAF, an advised fund of the Chan Zuckerberg Initiative Foundation.

## Notes

https://github.com/COMBINE-lab/simpleaf

https://github.com/COMBINE-lab/piscem

https://github.com/COMBINE-lab/alevin-fry/

https://github.com/COMBINE-lab/fast-flex-processing-paper-scripts

### Summary of Updates

This revision focuses on improving the writing clarity, exposition, and flow of the manuscript and supplement. The underlying methods, benchmarks, datasets, and quantitative results remain unchanged. Across the paper, we revised the prose to further emphasize intuition, to better defined specialized terminology at or before first use, and to tighten the narrative transitions. We also clarified memory comparisons by reporting relative peak usage directly as percentages alongside multiples, added brief discussion of architectural scope and assay boundaries, and updated section numbering and cross-references for consistency.

