## Supplementary Information for "Rapid, accurate and memory-frugal sub-gene-level quantification of probe-based, multiplexed single-cell data"


| Dataset | Download link |
|---|---|
| 128k human PBMC v1 16plex | https://www.10xgenomics.com/datasets/128k-human-pbmcs-stained-with-totalseqc-human-universal-cocktail |
| 320k human PBMC v1 16plex | https://www.10xgenomics.com/datasets/320k_Human_PBMCs_Sub_Pool_16-plex_GEM-X_FLEX |
| 40k human DTC v1 4plex | https://www.10xgenomics.com/datasets/40k-mixture-of-dissociated-tumor-cells-from-3-donors-stained-with-totalseqc-antibodies |
| 16k human PBMC v2 4plex | https://www.10xgenomics.com/datasets/Flexv2_16k_Human_PBMCs_TotalseqC_4plex |
| 16k mouse splenocyte v2 4plex | https://www.10xgenomics.com/datasets/Flexv2_16k_mouse_Splenocytes_TotalseqC_4plex |

Supp. Table 1: Dataset download links.

### S1 Sources of residual discordance between `simpleaf` and `Cell Ranger`

We analyzed the 16k mouse splenocyte Flex v2 library to identify the main sources of residual discordance between `simpleaf` and `Cell Ranger`. Out of 1,185,501,000 read pairs, `simpleaf` excluded 7.16% during R1 geometry parsing, 0.91% without an R2 probe placement, and 0.02% matching excluded-probe decoys (probes flagged as excluded in the panel specification and indexed as decoys so off-target reads are discarded instead of mis-mapping to included probes); overall, 91.90% of read pairs mapped and 87.50% remained after barcode correction and collation.

During R1 parsing, `simpleaf` retained reads with one anchor mismatch (1.67% of all read pairs) and reads with a four-nucleotide spacer (0.19%), while excluding reads with an anchor upstream of the configured window (4.60%), downstream of the window (0.42%), or lacking an anchor match within one substitution (2.14%) (Supp. Table 2).

`Cell Ranger` accepts several R1 geometries that `simpleaf` excludes. In particular, an anchor at zero-based position 27 (one nucleotide upstream of `simpleaf`'s window, which starts at position 28 immediately after the 16-base cell barcode and 12-base UMI) occurred in 1.64% of read pairs; at that offset, `Cell Ranger`'s fixed UMI interval overlaps the anchor by one base. These differing extraction rules account for part of the residual UMI count difference.

R2 mapping differences arise from the tools' distinct matching criteria. Among 1.06 billion reads assigned to a probe by `Cell Ranger`, 14,287,014 (1.3%) failed its initial two-half matching criterion but were recovered by edit-distance rescue and still assigned to the same gene by `simpleaf`. Conversely, only 169,238 reads (0.02%) satisfied `Cell Ranger`'s initial criterion

| R1 read class | Outcome | Read pairs (%) |
|---|---|---|
| Anchor at positions 28–32 with at most one mismatch | R1 pass | 1,100,582,138 (92.84) |
| Correctable cell barcode | Corrected | 21,539,640 (1.82) |
| Correctable sample barcode | Corrected | 21,483,701 (1.81) |
| Anchor before position 28 | Excluded | 54,520,790 (4.60) |
| Anchor after position 32 | Excluded | 4,930,698 (0.42) |
| No anchor within one mismatch | Excluded | 25,426,844 (2.14) |
| Uncorrectable sample barcode | Excluded | 16,831,803 (1.42) |
| Uncorrectable cell barcode | Excluded | 35,342,432 (2.98) |

Supp. Table 2: R1 geometry parsing and barcode-assignment outcomes for the 16k mouse splenocyte Flex v2 library. Percentages are computed over all 1,185,501,000 read pairs. Positions are zero-based. Geometry and barcode-assignment rows are not mutually exclusive.

| Read class | `Cell Ranger` | `simpleaf` | Read count |
|---|---|---|---|
| Fails `Cell Ranger`'s initial two-half criterion; both tools assign the same gene | Assigned after rescue | Same-gene assignment | 14,287,014 |
| Passes `Cell Ranger`'s initial two-half criterion; `simpleaf` does not assign the same gene | Assigned | No same-gene assignment | 169,238 |

Supp. Table 3: R2 mapping classifications among 1.06 billion `Cell Ranger` probe-assigned reads in the 16k mouse splenocyte Flex v2 library. `simpleaf` used a probe index with $k = 19$.

but were not assigned to that gene by `simpleaf` (Supp. Table 3). Gene reassignment affected 29,030 molecules (0.011%).

Across the four mouse samples, `simpleaf` achieved a mean count concordance with `Cell Ranger` of 98.61% and a mean per-gene Spearman correlation of 0.9951. Across all five libraries, mean per-cell Spearman correlations ranged from 0.966 to 0.997 and count concordance ranged from 97.8% to 99.6%.

### S2 Implementation details for the computational approach

Here we detail the storage layouts, memory accounting, and execution paths of the lookup, barcode-correction, and collation stages introduced in Section 2.

**S2.1 Probe-panel dictionary characterization.** We characterized the three probe-panel indexes using `simpleaf` commit `645841e` and `piscem` commit `b19ee69` (`piscem-rs` 0.11.1 and `tiny-dict` 0.8.0), with $k = 19$, minimizer length $m = 15$, and excluded probes retained as poison decoys. The probe CSVs were obtained from the pinned chemistry registry, with matching BLAKE3 hashes, and measured on an AMD EPYC 9575F host (Supp. Table 4).

Using a read-only diagnostic, we loaded each index through the mapper's loader and tallied requested heap allocations for the hash-table buckets and control bytes, packed reference strings and offsets, segment-occurrence arrays and overflow lists, and the 2 MiB load-time Bloom prefilter. Component sums were cross-checked against live requested-byte deltas from a counting wrapper around mimalloc after temporary loading allocations were freed. These totals represent requested heap bytes for lookup structures rather than whole-index memory or peak process RSS, and exclude shallow struct headers, allocator metadata and size-class rounding, reference-name tables, equivalence-class and poison tables, and read-processing buffers. Inline percentages are computed over all segments without query-frequency weighting; runtime miss-rate and filter-activation statistics were not recorded.

**S2.2 Packed index and barcode schema.** Each hash-table entry consists of a 64-bit canonical $k$-mer key alongside a 64-bit value containing a segment identifier, within-segment offset, and canonical-orientation flag. Co-locating the key and value avoids a separate value-array lookup and lets the mapper verify exact $k$-mer membership without decoding reference

| Probe panel | Probes | Canonical keys | Segments | Inline (%) | Lookup (MiB) |
|---|---|---|---|---|---|
| Human v1.1.0 | 53,459 | 1,710,227 | 53,740 | 99.78 | 37.46 |
| Human v2.0.0 | 53,519 | 1,712,147 | 53,800 | 99.78 | 37.46 |
| Mouse v2.0.0 | 55,178 | 1,765,030 | 55,691 | 99.63 | 37.52 |

Supp. Table 4: Characterization of the probe-panel indexes. Lookup allocations cover the hash table, packed strings and offsets, occurrence storage, and a 2 MiB prefilter; MiB denotes $2^{20}$ bytes. Inline is the percentage of segments with a single probe occurrence stored inline. Lookup allocations denote requested heap bytes rather than peak process RSS.

strings. The mapper then resolves probe coordinates in the segment-occurrence table. For single-probe segments, the probe identifier, offset, and orientation fit inline in one 64-bit entry, whereas entries for multi-probe segments contain an overflow-list index, so single-probe hits require no secondary list fetch.

In RAD v2, hierarchical barcoding is recorded once in the file prelude's read-tag schema (Supp. Figure 1). Each barcode tag has an integer storage type and a `Barcode` role with a hierarchy level and nucleotide length (level 0 for the sample barcode and level 1 for the cell barcode in Flex), while the UMI length is recorded in a `Umi` role. Each read record then contains its alignment count, the two-bit-encoded barcode and UMI integers in schema order, and the probe mapping assignments. Downstream tools reconstruct the composite sample–cell key directly from these tag roles, so identical cell-barcode sequences from different samples stay in separate UMI-resolution groups.

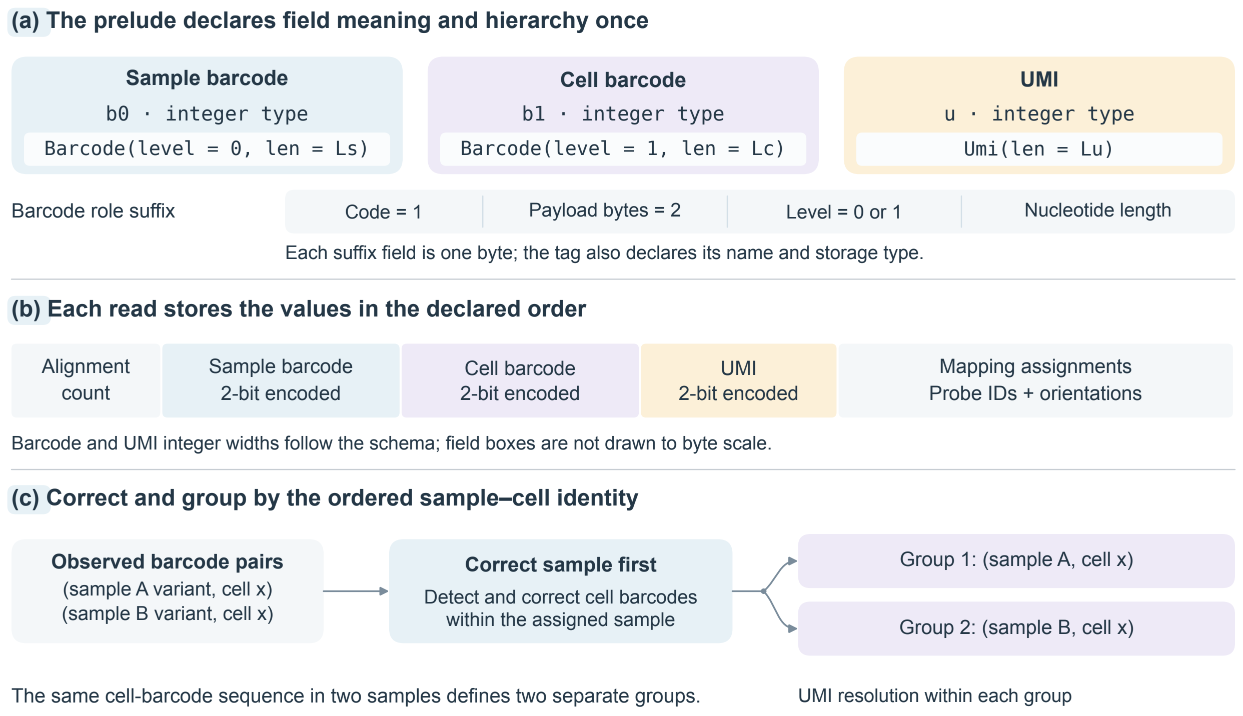


Supp. Figure 1: Encoding and processing hierarchical barcodes in RAD v2. (a) Field types and semantic roles are stored once in the file prelude as read-tag descriptors, including the hierarchy level and nucleotide length for `Barcode` roles and the nucleotide length for the `Umi` role. Each role suffix contains a code, a payload length, and the role-specific payload. (b) Each read record contains an alignment count, the encoded sample and cell barcodes, the UMI, and mapping assignments. (c) Sample-barcode correction precedes cell-barcode detection and correction within each sample. Collation groups records by the corrected sample–cell pair, keeping identical cell-barcode sequences from different samples separate. Sample and cell symbols are schematic; no experimental counts are shown.

#### S2.3 Permit-list generation and correction plans.

During permit-list generation, `alevin-fry` routes reads with unambiguous sample barcodes through a precomputed lookup table. Worker threads accumulate cell-barcode counts in bounded local histograms and periodically merge them into per-sample histograms; this reduces synchronization overhead and stores one count per distinct barcode instead of individual raw occurrences. Within each sample, permitted barcodes are stored as sorted, two-bit-encoded integers with a prefix-offset directory so candidate lookups binary-search only a single prefix slice, and one-substitution neighbors are generated directly on the packed integers. Exact matches take precedence, while ambiguous one-mismatch candidates are resolved under the configured unique-target or frequency-based policy using priors fixed from exact-match counts. `alevin-fry` serializes one decision per distinct observed cell barcode in a sample-scoped correction plan for reuse during collation. Samples are processed in parallel up to the thread budget and estimated per-sample memory limits.

Before collation, `alevin-fry` assigns corrected sample–cell groups to logical buckets from their read counts without splitting any group, and compiles the correction plan into a sample-routing table and per-sample cell lookup tables. Given an input sample–cell pair, sample routing followed by one cell lookup returns both the corrected cell barcode and its bucket identifier, packed into a single 64-bit integer when narrow enough or stored in a wider entry otherwise. Neither neighbor generation nor a separate bucket lookup is repeated per read during scatter.

#### S2.4 Scatter, gather, and memory controls.

Scatter workers read RAD chunks against this immutable shared plan, buffer records by logical bucket, flush them to one private, append-only spool file per worker, and log the byte extents of each bucket. Physical temporary-file count therefore equals the scatter worker count rather than the potentially much larger number of logical buckets. Gather workers then replay each bucket's extents in two linear passes. In the first pass, a worker reads record headers to compute per-group byte totals and prefix-summed output offsets while saving a 32-bit group index per record. In the second pass, it uses that index to copy raw record bytes into each group's contiguous output region without re-hashing composite keys, running a comparison sort, or decoding alignment payloads.

Spool-buffer sizes and concurrent gather workers are bounded from bucket-size estimates and a user-overridable working-memory target, and `alevin-fry` frees the scatter lookup tables before allocating gather output buffers so the two working sets do not overlap.

### S3 Additional concordance and probe-level analyses

Clustering and marker-gene agreement across all five benchmarked libraries are summarized in Supp. Table 5; equal cluster counts were obtained in all 132 pairwise comparisons.

When comparing UMI-resolution scopes, summed per-probe totals exceeded direct gene-level UMI totals by 0.0036–0.2257% under one-mismatch UMI correction, because one-mismatch UMIs on different probes of the same gene can be collapsed together during gene-level resolution while remaining separate per probe. Without UMI error correction, summed probe-level totals were 0.00068–0.0167% lower than gene-level totals, because UMIs with tied read support across sibling probes remain assignable to the gene as a whole but are discarded as ambiguous at the probe level.

Cell-type annotations and cell counts for the 320k human PBMC UMAP embedding in Figure 4 are shown in Supp. Figure 3.

| Dataset | Homogeneity | Completeness | ARI | Marker Jaccard | Shared top-50 markers |
|---|---|---|---|---|---|
| 16k human PBMC v2 4plex | 0.936 | 0.943 | 0.873 | 0.869 | 45.9 |
| 16k mouse splenocyte v2 4plex | 0.876 | 0.874 | 0.795 | 0.780 | 42.5 |
| 40k human DTC v1 4plex | 0.781 | 0.784 | 0.631 | 0.682 | 38.4 |
| 128k human PBMC v1 16plex | 0.936 | 0.938 | 0.890 | 0.852 | 45.0 |
| 320k human PBMC v1 16plex | 0.869 | 0.864 | 0.720 | 0.838 | 44.4 |

Supp. Table 5: Clustering and marker-gene agreement between `simpleaf`, `cyto` and `Cell Ranger` across the five datasets. Clustering agreement is summarized by homogeneity, completeness and adjusted Rand index (ARI). Marker-gene overlap is summarized by the mean Jaccard index and the mean number of shared genes among the top 50 markers per cluster.

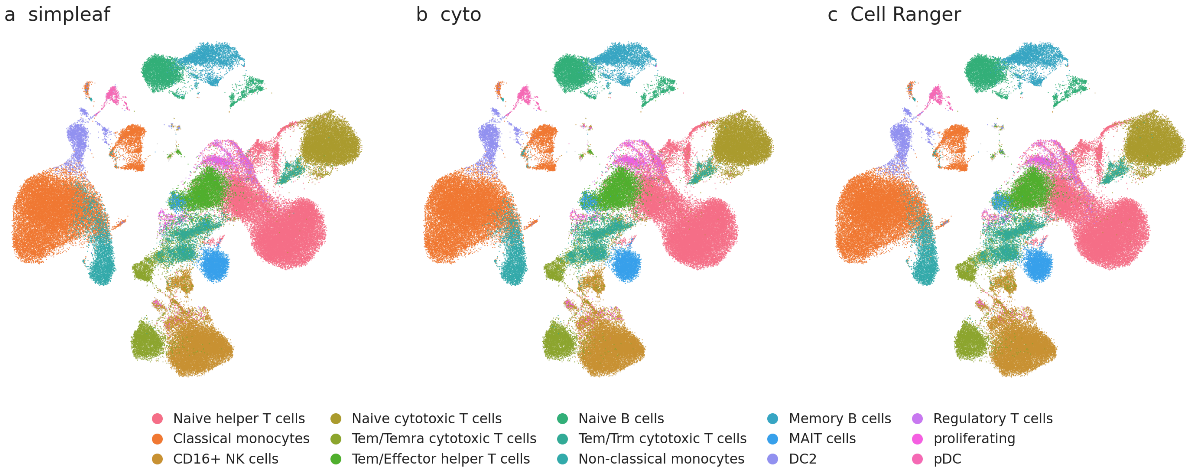


Supp. Figure 2: UMAP embedding of the 16 pooled samples of the 128k human PBMC library, computed from the `Cell Ranger` count matrix using the workflow described in Section 5.4. The analysis included 121,652 `Cell Ranger`-called cells that passed quality control in all three tools. All three panels show the same embedding, and each cell is colored by the cell type assigned from the `simpleaf` (a), `cyto` (b), and `Cell Ranger` (c) counts, using the cell type annotation described in Section 5.4.5. Of these cells, 93.9% have the same cell type from `simpleaf` as from `Cell Ranger`, and 94.4% from `cyto`. No batch correction was applied.

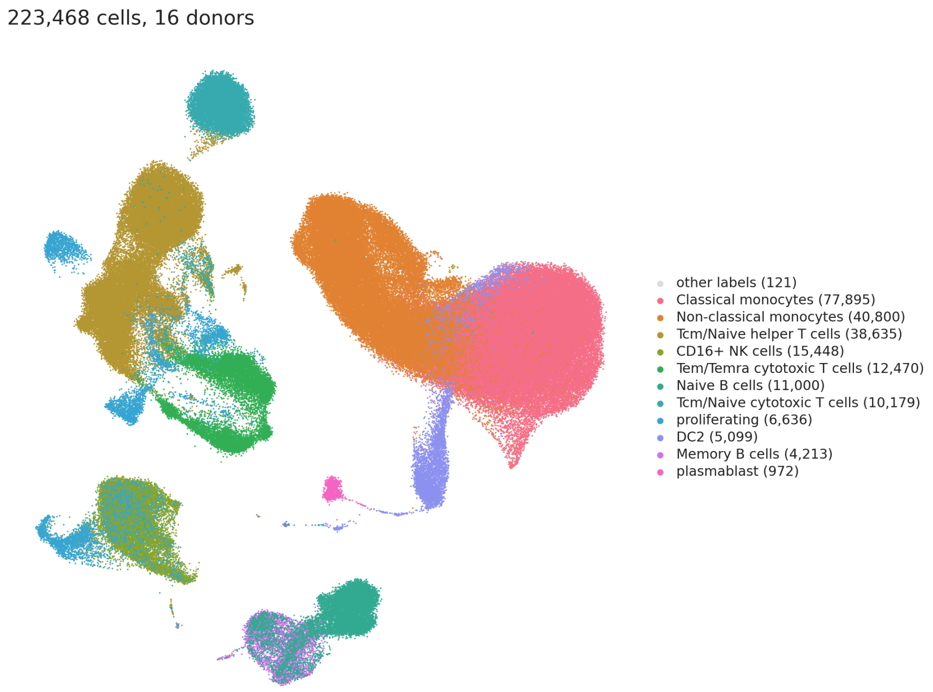


Supp. Figure 3: Cell-type annotations and cell counts for the UMAP embedding in Figure 4. The embedding includes 223,468 cells from all 16 donors after doublet removal.
